# Metabolic modelling of the C_3_-CAM continuum revealed the establishment of a starch/sugar-malate cycle in CAM evolution

**DOI:** 10.1101/2019.12.16.877621

**Authors:** Ignacius Y. Y. Tay, Kristoforus Bryant Odang, C. Y. Maurice Cheung

## Abstract

The evolution of Crassulacean acid metabolism (CAM) is thought to be along a C_3_-CAM continuum including multiple variations of CAM such as CAM cycling and CAM idling. Here, we applied large-scale constraint-based modelling to investigate the metabolism and energetics of plants operating in C_3_, CAM, CAM cycling and CAM idling. Our modelling results suggested that CAM cycling and CAM idling could be potential evolutionary intermediates in CAM evolution by establishing a starch/sugar-malate cycle. Our model analysis showed that by varying CO_2_ exchange during the light period, as a proxy of stomatal conductance, there exists a C_3_-CAM continuum with gradual metabolic changes, supporting the notion that evolution of CAM from C_3_ could occur solely through incremental changes in metabolic fluxes. Along the C_3_-CAM continuum, our model predicted changes in metabolic fluxes not only through the starch/sugar-malate cycle that is involved in CAM photosynthetic CO_2_ fixation but also other metabolic processes including the mitochondrial electron transport chain and the tricarboxylate acid cycle at night. These predictions could guide engineering efforts in introducing CAM into C_3_ crops for improved water use efficiency.

## Introduction

Crassulacean acid metabolism (CAM) photosynthetic CO_2_ fixation is an evolutionary descendant of C_3_ photosynthesis, which is known to have evolved independently multiple times in at least 35 plant families comprising about 6% of flowering plant species (Winter and Smith, 1996a; Silvera et al., 2010). CAM is an adaptation of photosynthetic CO_2_ fixation typically associated to limited water availability (Cushman and Borland, 2002). By closing their stomata during the light period and fixing atmospheric and/or respiratory carbon dioxide (CO_2_) exclusively in the dark period, CAM allows plants to use water more efficiently while fixing carbon for growth. The engineering of CAM into C_3_ crops has been suggested as a possible strategy to meet the demands on agriculture for food, feed, fibre, and fuels, without exacerbating the pressures on arable land area due to climate change (Borland et al., 2014). However, as a carbon-concentrating mechanism, CAM is thought to be more metabolically expensive than C_3_ (Winter and Smith, 1996b), which suggests that transferring a CAM pathway into C_3_ crops would incur a crop yield penalty. To investigate the energetics of C_3_ and CAM, large-scale metabolic models were applied which showed that engineering CAM into C_3_ plants does not impose a significant energetic penalty given the reduction in photorespiration from the carbon-concentrating mechanism (Cheung et al., 2014; Shameer et al., 2018).

Besides the phylogenetic and ecological diversity of CAM plants, there is remarkable plasticity in its metabolism with multiple defined variations of CAM including CAM cycling and CAM idling (Ting, 1985; Cushman, 2001, Winter, 2019). Briefly, CAM cycling primarily fixes CO_2_ in the light period with refixing respiratory CO_2_ behind closed stomata at night, leading to a small diel organic acid flux; CAM idling lacks diel gaseous exchange with closed stomata across the 24 hour light/dark cycle and has a small continued diel fluctuation in organic acids level (Sipes and Ting, 1985; Cushman, 2001, Winter, 2019). Silvera et al. (2010) generalised the idea of plasticity of CAM into a continuum of CAM levels, due to the differences in the degree of nocturnal and daytime net CO_2_ uptake. Bräutigam et al. (2017) took the idea further to include C_3_ as part of the C_3_-CAM continuum and suggested that the evolution of C_3_ to CAM only required incremental increases in metabolic fluxes. In this study, large-scale metabolic modelling was applied to investigate how CAM cycling and CAM idling fit into the continuum of CAM evolution and to identify the changes in metabolic fluxes along the C_3_-CAM continuum. The results from our modelling study provide novel insights into the energetics and metabolic alterations from C_3_ to CAM, which could guide engineering efforts aimed at introducing CAM into C_3_ plants.

## Materials and Methods

### Core metabolic model for modelling C_3_, CAM, CAM cycling and CAM idling

The mass- and charge-balance core metabolic model of Arabidopsis in Shameer et al. (2018) was used for modelling the metabolism of leaves operating in C_3_, CAM, CAM cycling and CAM idling. A number of minor modifications were made to the Shameer et al. (2018) model to more accurately model metabolism of C_3_, CAM, CAM cycling, CAM idling and the C_3_-CAM continuum. Firstly, a reaction for the accumulation of oxygen, which is produced from water splitting in the photosynthetic light reactions, was added to the model such that we could run the simulations for CAM and CAM idling as oxygen exchange was constrained to zero during the day in these two scenarios. Another modification from Shameer et al. (2018) was how the acidification of the vacuole was modelled. Instead of directly setting different fixed pH for the vacuole in C_3_ and CAM, protons were allowed to accumulate in the vacuole in our model. A reaction allowing protons to freely flow in and out of the cytosol was blocked such that pH homeostasis can be modelled through the accumulation of protons in the vacuole. For linking the cytosolic and mitochondrial proton pools, proton transport between the cytosol and the mitochondrial intermembrane space was set to be reversible. The modification of this constraint only led to a very minor change in the flux predictions (data not shown). Irreversible proton transporters were added from the vacuole to the cytosol and from extracellular to the cytosol to allow leakage of protons down the electrochemical gradients. Lastly, the compartmentation of metabolites in the reaction “HEXOKINASE_RXN_MANNOSE_c” was corrected to be in the cytosol. The modified core model can be found in SBML and Excel formats (Supplementary File S1).

### Model simulations with flux balance analysis

Based on the constraints and objective function stated in the Results section, parsimonious flux balance analysis (pFBA) was performed using scobra (https://github.com/mauriceccy/scobra), an extension of cobrapy (Ebrahim et al., 2013). The scripts for running the simulations in this study can be found in Supplementary File S2. In this study, we primarily reported the results from the pFBA simulations (Supplementary Tables S1, S2, S3). The conclusions made based on the pFBA results for C_3_, CAM, CAM cycling and CAM idling were confirmed using flux variability analysis (Mahadevan and Schilling, 2003) applied on the primary objective (Supplementary Table S4).

## Results

### Predicted metabolic fluxes of C_3_, CAM, CAM cycling and CAM idling

In this study, we simulated the metabolism of leaves undergoing C_3_, CAM, CAM cycling and CAM idling using a recently published core metabolic model of Arabidopsis which was used to model C_3_ and CAM plants (Shameer et al., 2018). Minor modifications of the model were outlined in the Materials and Methods section. The constraints for simulating the core metabolic functions of mature leaves, namely export of sucrose and amino acids into the phloem and cellular maintenance, were set based on the values in Shameer et al. (2018). All simulations, except CAM idling, were constrained to have a phloem export rate of 0.259 μmol m^−2^ s^−1^ based on the value of C_3_ plants in Shameer et al. (2018). The set of constraints for modelling the four different modes of photosynthesis are summarised in Table 1. The primary objective function of minimising photon demand was used throughout this study, which allows us to study the metabolic efficiencies of the different modes of photosynthesis. Parsimonious flux balance analysis (pFBA), i.e. minimisation of absolute sum of fluxes, was applied as a secondary objective to eliminate substrate cycles. Results from pFBA were confirmed using flux variability analysis (Mahadevan and Schilling, 2003) performed on the primary objective.

**Table 1:** Sets of constraints for modelling C_3_, CAM, CAM cycling and CAM idling. Phloem export rate was set based on the predicted value of C_3_ plants in Shameer et al. (2018). RuBisCO carboxylase:oxygenase ratio was set to 3:1 when stomata is opened, and 5.15:1 when stomata is closed based on Shameer et al. (2018).

| **Constraints**<br>($\mu\text{mol m}^{-2} \text{s}^{-1}$) | **C<sub>3</sub>** | **CAM** | **CAM Cycling** | **CAM Idling** |
| --- | --- | --- | --- | --- |
| Phloem export | 0.259 | 0.259 | 0.259 | 0 |
| CO<sub>2</sub> exchange (light) | Unconstrained | 0 | Unconstrained | 0 |
| CO<sub>2</sub> exchange (dark) | Unconstrained | Unconstrained | 0 | 0 |
| O<sub>2</sub> exchange (light) | Unconstrained | 0 | Unconstrained | 0 |
| O<sub>2</sub> exchange (dark) | Unconstrained | Unconstrained | 0 | 0 |
| RuBisCO<br>carboxylase:oxygenase<br>ratio (light) | 3:1 | 5.15:1 | 3:1 | 5.15:1 |
| RuBisCO<br>carboxylase:oxygenase<br>ratio (dark) | 3:1 | 3:1 | 5.15:1 | 5.15:1 |

The model predictions of C_3_ and CAM were very similar to that in Shameer et al. (2018) given the similarities in the constraints used. Without any constraints on malate decarboxylation enzyme and carbohydrate storage, the model predicted net carbon fixation during the light period in the C_3_ flux prediction, whereas in CAM carbon was fixed in the dark period with phospho*enol*pyruvate carboxykinase (PEPCK) being the main predicted route for malate decarboxylation. Starch was predicted to be the main carbohydrate storage in both C_3_ and CAM. These results are consistent with the findings in Shameer et al. (2018) where starch-storing PEPCK subtype were predicted to be the most energy efficient. The effect of the choice of decarboxylation enzymes (PEPCK vs malic enzyme) on the model predictions was explored by constraining other decarboxylating enzymes to carry zero flux. It was found that the choice of decarboxylation enzymes makes little qualitative difference with respect to the results presented (Supplementary Table S5). From here on, the results presented were model predictions with no constraints on the decarboxylation enzymes. As for carbohydrate storage, simulations were performed with starch, sucrose or fructan as the sole carbohydrate storage. Except for reactions involved in the synthesis, accumulation and degradation of carbohydrate storage, the predicted fluxes in central carbon metabolism were largely similar between the three carbohydrate storages tested (Supplementary Table S6). In this study, we mostly presented the results from simulations with starch as the carbohydrate storage. Similar conclusions can be made for using sugar as the carbohydrate storage. The core set of metabolic fluxes for C_3_, CAM, CAM cycling and CAM idling with starch as the carbohydrate storage is depicted in Figure 1.

**Figure 1.**
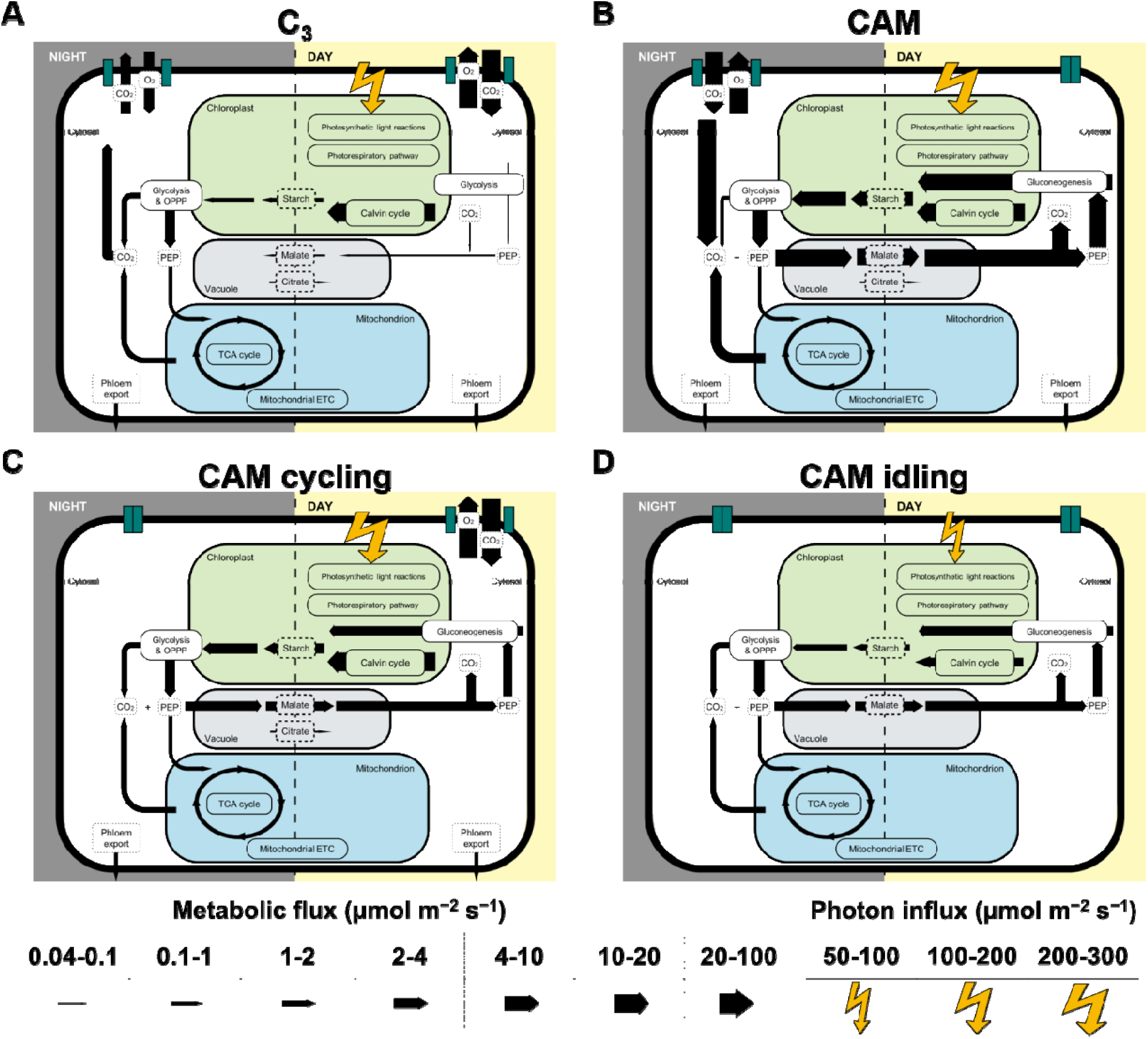
Core sets of metabolic fluxes in the four modes of photosynthesis modelled: (A) C_3_, (B) CAM, (C) CAM cycling and (D) CAM idling. The width of the arrows represents the magnitude of the reaction flux according to the scale on the bottom of the figure in μmol m^−2^ s^−1^. The photorespiratory pathway is shown in chloroplast for simplicity, which in reality spans multiple compartments. Flux from 3-phosphoglycerate to PEP was taken as the flux for glycolysis and gluconeogenesis. Flux for succinate dehydrogenase was taken as the TCA cycle flux. RuBisCO carboxylase flux was taken as the flux through the Calvin-Benson cycle.

#### CAM cycling

Similar to C_3_ plants, CAM cycling fixes carbon in the light period. CAM cycling is characterised by its closed stomata in the dark period with refixation of respiratory CO_2_ and a small diel organic acid flux (Sipes and Ting, 1985; Cushman, 2001, Winter, 2019). To model CAM cycling, we applied the C_3_ constraints with an additional constraint of setting CO_2_ and O_2_ exchange at night to zero to simulate the closure of the stomata (Table 1). This resulted in a flux distribution that resembled a weak version of CAM, with nocturnal malate accumulation and increased light period starch accumulation (Figure 1C). Phospho*enol*pyruvate carboxylase (PEPC) was predicted to be active only at night in CAM cycling for CO_2_ refixation, in contrast to C_3_ where PEPC was only active during the light period (Supplementary Table S2). Another major difference between CAM cycling and C_3_ is malate accumulation. While C_3_ was predicted to have a very small amount of malate accumulation during the light period, CAM cycling was predicted to have substantial amount of nocturnal malate accumulation (∼20% of the amount of malate accumulation in CAM) (Figure 1; Supplementary Table S2), which is consistent with known behaviour of CAM cycling (Ting, 1985; Cushman, 2001). The nocturnal malate accumulation and respiratory CO_2_ refixation via PEPC under the CAM cycling scenario were accompanied by changes in fluxes in other parts of metabolism. Malate decarboxylation during the light period was predicted to be active in CAM cycling but not in C_3_ (Figure 1). There was a larger flux through gluconeogenesis to convert malate into starch in the light period, which led to more starch accumulation during the light period in CAM cycling compared to C_3_ (Figure 1; Supplementary Table S2). Given that CAM cycling has a higher starch accumulation in the light period, it was predicted to have a larger glycolytic flux in the dark to convert starch into phospho*enol*pyruvate (PEP) for CO_2_ refixation, compared to C_3_ (Figure 1; Supplementary Table S2). The activities of most of the other reactions at night were similar in CAM cycling and in C_3,_ with CAM cycling having a slightly higher flux through the tricarboxylic acid (TCA) cycle and the mitochondrial electron transport chain (ETC), presumably to produce extra ATP for transporting malate into the vacuole for storage at night.

#### CAM idling

CAM idling is characterised by the lack of diel gaseous exchange and a small continued diel fluctuation in the organic acids level because of internally recycled CO_2_ (Sipes and Ting, 1985, Winter, 2019). It is usually an adaptation in water-stressed plants, which results in the closure of stomata for the whole 24-hour cycle. To model this, the CO_2_ and O_2_ exchange during the light and the dark periods were constrained to carry zero flux (Table 1). Given that there is no CO_2_ exchange, we assumed that there is no net carbon fixation, hence phloem export was constrained to zero for CAM idling.

The primary metabolic demand for plants in CAM idling is cellular maintenance. The model predicted a starch-malate cycle where starch accumulated in the light period is metabolised in the dark period mainly through glycolysis and the oxidative pentose phosphate pathway (OPPP) to produce ATP and NADPH for maintenance processes (Figure 1D). While the majority of PEP was used as precursor for carbon refixation by PEPC, a significant proportion of PEP was predicted to be metabolised further through the TCA cycle to feed the mitochondrial ETC for ATP synthesis (Figure 1D). Given that it is a closed system with respect to carbon, CO_2_ produced in the OPPP and the TCA cycle is refixed by PEPC, which ultimately leads to the accumulation of malate in the dark. In the light period, PEP from malate decarboxylation was recycled to produce starch via gluconeogenesis, while the CO_2_ produced from malate decarboxylation was refixed via the Calvin-Benson cycle similar to the scenario for CAM (Figure 1). With no net carbon import or export, the amount of carbon stored in starch in the light period was predicted to be equalled to the amount of carbon storage in malate at night. The starch-malate cycle was primarily driven by the energy from the light reactions of photosynthesis, and it acted as a carbon neutral way of storing and transferring energy from the light period to the dark period. Similar results were obtained when sucrose or fructan was used as the sole carbohydrate storage instead of starch (Supplementary Table S6), meaning that a sugar-malate cycle can serve the same function as the starch-malate cycle in sugar-storing plants.

### Energetics and metabolite accumulation in C_3_, CAM, CAM cycling and CAM idling

The metabolic flux predictions of C_3_, CAM, CAM cycling and CAM idling were compared to see how CAM cycling and CAM idling fit into the evolution of CAM from C_3_. Table 2 summarises the predicted fluxes related to energetics and metabolic accumulation in the four simulations. CAM idling was predicted to use the fewest photons, which was expected given that it does not have the metabolic demand for exporting sucrose and amino acids into the phloem. For the same metabolic demand, CAM requires more photons than C_3,_ as expected. It is interesting to see that the photon demand for CAM cycling falls between C_3_ and CAM. A similar trend was observed for other fluxes related to energy metabolism including the ATP and NADPH production by the photosynthetic light reactions and the ATP production by the mitochondrial ATP synthase (Table 2). The same trend was also reflected in the energetic demands of the Calvin-Benson cycle in terms of ATP and NADPH consumption (Supplementary Table S2).

**Table 2.**
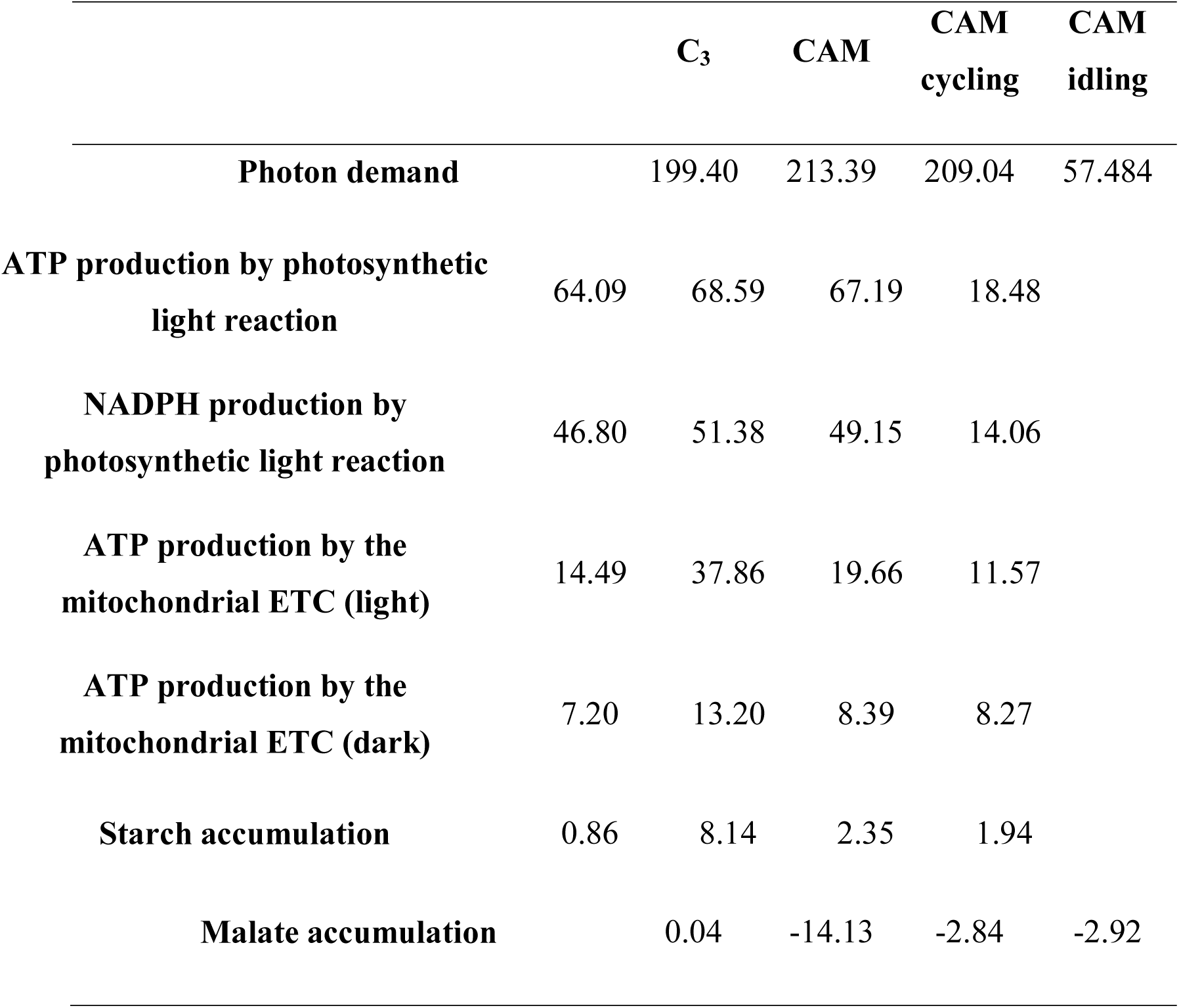
Fluxes related to energetics and metabolic accumulation predicted in the model simulations of C_3_, CAM cycling, CAM idling and CAM. Photon demand and the productions of ATP and NADPH by photosynthetic light reaction are flux values from the light period. A positive value of metabolite accumulation denotes a net accumulation in the light period; negative value of metabolite accumulation denotes a net accumulation in the dark period. All values are in the units of μmol m^−2^ s^−1^.

Metabolite accumulation showed a different pattern compared to the energetics (Table 2). C_3_ had the lowest daytime starch accumulation, followed by CAM cycling and CAM idling which had about 2-3 times more starch accumulation than C_3_. CAM had the highest light period starch accumulation with more than nine times the amount associated with C_3_. This suggested that CAM cycling and CAM idling could potentially be intermediate steps in CAM evolution with respect to the regulation of starch accumulation. A similar pattern can be observed for malate accumulation. A very small amount of malate was predicted to accumulate during the day for C_3_ plants, whereas a large nocturnal malate accumulation was predicted for CAM as part of CAM photosynthesis. CAM cycling and CAM idling had intermediate level of nocturnal malate accumulation (∼20% of that in CAM), which was related to the refixation of nocturnal CO_2_ by PEPC. Reactions related to the starch/sugar-malate cycle, including glycolysis and PEPC flux in the dark period, and gluconeogenesis and malate decarboxylation during the light period, showed a similar trend (Supplementary Table S2) suggesting that CAM cycling and/or CAM idling could be an evolutionary intermediate for the evolution of the extensive starch/sugar-malate cycle in CAM plants.

### Predicting the metabolic transitions during C_3_-CAM evolution

The behaviour of diel CO_2_ exchange is the main diagnostic indicator between C_3_ and CAM (Silvera et al., 2010). To model the potential metabolic transitions that could happen during the evolution of CAM from C_3_, we varied the CO_2_ uptake rate during the light period from 13.12 μmol m^−2^ s^−1^ (the predicted value for C_3_) to 0 μmol m^−2^ s^−1^ (which had the same effect as gradually increasing nocturnal CO_2_ uptake given the overall carbon balance). This simulates the decrease in gaseous exchange during the light period by stomatal closure, hence a similar constraint was set for light period oxygen exchange. As the stomata closes in the light period, i.e. light period CO_2_ uptake decreases, it was assumed that the proportion of ribulose-1,5-bisphosphate carboxylase/oxygenase (RuBisCO) flux going through the carboxylase reaction increases linearly from 75% (carboxylase to oxygenase ratio of 3:1) to 83.74% (carboxylase to oxygenase ratio of 5.15:1) to account for the reduction of photorespiration. All other constraints remained the same as the C_3_ and CAM simulations. This analysis simulates the closing of stomata which decreases atmospheric CO_2_ intake during the light period. The full results from this simulation can be found in Supplementary Table S3.

Given that the metabolic demands remained constant throughout the analysis, a decrease in CO_2_ uptake in the light period led to a shift from C_3_ to CAM photosynthesis with an increase in flux through the starch-malate cycle including starch degradation, glycolysis, PEPC, and malate accumulation at night, and malate decarboxylation and starch accumulation during the light period (Figure 2A,B; Supplementary Table S3). Note that dark period CO_2_ uptake increased as light period CO_2_ uptake decreased due to the carbon balance of the model in exporting a fixed amount of sucrose and amino acids into the phloem. CAM cycling occurs at the point when dark period CO_2_ uptake is zero.

**Figure 2.**
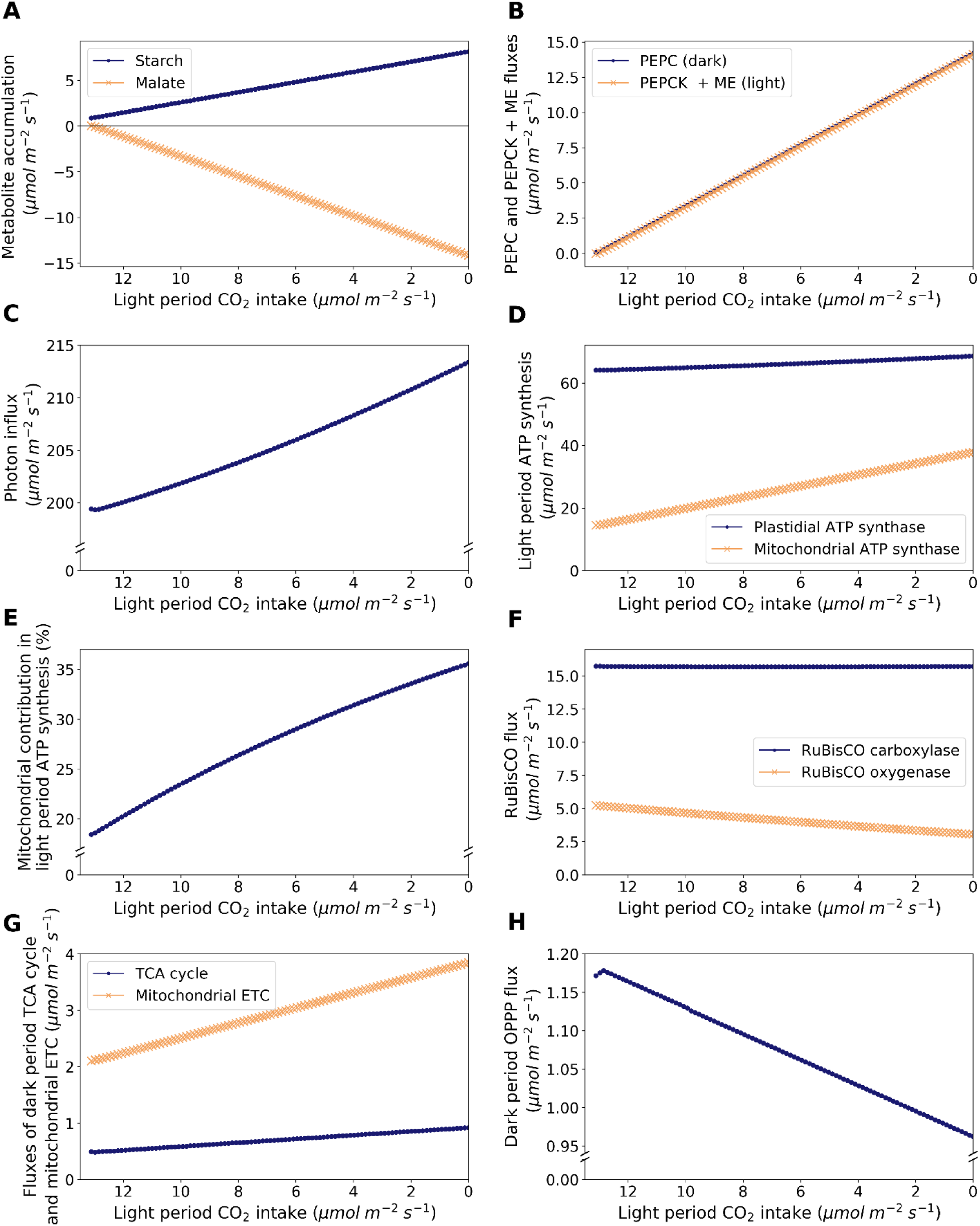
Model predictions of metabolic changes along the C_3_-CAM continuum, as modelled by varying CO_2_ exchange during the light period. (A) Accumulation of starch (dots) and malate (crosses), (B) Dark period PEPC flux in the dark period (dots) and malate carboxylation flux as the sum of fluxes of PEPCK and malic enzyme in the light period (crosses), (C) Photon intake in the light period, (D) ATP synthesis in the light period by plastidial ATP synthase (dots) and mitochondrial ATP synthase (crosses), (E) Proportion of light period ATP synthesis by the mitochondrial ATP synthase, (F) Fluxes of RuBisCO carboxylase (dots) and oxygenase (crosses), (G) Fluxes through the TCA cycle (taken as the flux of succinate dehydrogenase; dots) and the mitochondrial ETC (taken as the flux of NADH dehydrogenase; crosses) in the dark period, and (H) flux through the OPPP (taken as the sum of fluxes of plastidial and cytosolic glucose 6-phosphate dehydrogenases) in the dark period.

Despite the constrained decrease in RuBisCO oxygenase contribution as light period CO_2_ uptake decreased, the amount of energy (in terms of photons) required to sustain the same metabolic demand increased by about 7% from C_3_ to CAM (Figure 2C) as extra energy is needed to run the starch-malate cycle. This is correlated with the increase in flux through the photosynthetic light reactions. Besides plastidial ATP synthesis, there was also an increase in ATP synthesis by the mitochondrial ETC in the light period as the simulation shifted from C_3_ to CAM (Figure 2D). The contribution of mitochondrial ATP synthesis increased from 18.2% in C_3_ to 35.6% in CAM (Figure 2E), which is likely to be related to the increase in NADH produced during malate decarboxylation. In our simulations, the RuBisCO carboxylase flux was predicted to be remain relatively constant while the total RuBisCO flux (carboxylase + oxygenase) decreased from C_3_ to CAM due to the decrease in RuBisCO oxygenase activity (Figure 2F). There were two major factors affecting RuBisCO carboxylase flux, i) refixation of photorespiratory CO_2_, and ii) starch accumulation to support energy demand in the dark period. In this case, the two factors counteract each other throughout the simulation where photorespiration decreases and the energy demand for running the starch-malate cycle (mostly for pumping malate into the vacuole) increases from C_3_ to CAM. For the simulations with sucrose or fructan as the sole carbohydrate storage, the model predicted an increase in RuBisCO carboxylase flux from C_3_ to CAM as the energy required for running the sugar-malate cycle is higher than the starch-malate cycle (due to the cost of pumping sugars into the vacuole for storage).

During the night, other than the increase in glycolytic flux as part of the starch-malate cycle from C_3_ to CAM, the model predicted an 87% increase in flux through the TCA cycle and an 83% increase in flux through the mitochondrial ETC (Figure 2G). This increase in mitochondrial ATP synthesis was mostly used to support the ATP-dependent tonoplast proton pump for the increasing nocturnal vacuolar malate accumulation. The cytosolic OPPP flux was predicted to decrease by 30% in the night from C_3_ to CAM (Figure 2h). This could be explained by the increase in the TCA cycle flux which contributed to the production of NADPH in the mitochondrion by the NADP-isocitrate dehydrogenase. This lessened the demand for the production of cytosolic NADPH required to be shuttled into the mitochondrion for maintenance processes.

## Discussion

### CAM cycling and CAM idling as viable evolutionary steps for establishing the starch-malate cycle

CAM cycling is considered as a weak form of CAM with stomata are open during the day and are closed at night (Lüttge, 2004; Silvera et al., 2010, Winter, 2019). With these constraints, our model predicted the known features of CAM cycling including the refixation of respiratory CO_2_ in the dark period, and a small amount of nocturnal malate accumulation (Cushman, 2001, Winter, 2019). To support these metabolic behaviours, our model predicted the establishment of a starch-malate cycle in CAM cycling, which included increased flux through malate decarboxylation, gluconeogenesis and starch synthesis and accumulation during the light period, and starch degradation and glycolysis during the dark period, when compared to C_3_ plants. The main metabolic advantage of CAM cycling over C_3_ is its higher carbon conversion efficiency when photosynthesis is limited by stomatal conductance in the light period, i.e. carbon limited. Given the same metabolic outputs, CAM cycling was predicted to require 20% less external CO_2_ compared to C_3_ due to the refixation of nocturnal respiratory CO_2_. This comes with a minor cost of 4.8% more photons and 1.6% more RuBisCO activity required, assuming that there is no reduction in photorespiration, which could be affected by limiting stomatal conductance and internal CO_2_ generation from malate decarboxylation. Given an environment that limits stomatal conductance in the light period, e.g. high temperature and drought, the evolution of CAM cycling, together with the establishment of the starch/sugar-malate cycle, was predicted to be advantageous in maximising carbon conversion efficiency. The metabolic activities of all reactions in the starch-malate cycle in CAM cycling were predicted to be at an intermediate level between C_3_ and CAM. The same applies to other supporting reactions such as the TCA cycle in the dark and the mitochondrial ETC during the light and dark periods. These findings suggest that CAM cycling is likely to be a possible evolutionary step along the path to the evolution of CAM.

As opposed to CAM cycling, CAM idling is thought of as a form of very strong CAM (Lüttge, 2004, Winter, 2019). In CAM idling, stomata remain closed throughout the day and night with small, sustained diel fluctuations in organic acids (Cushman, 2001; Silvera et al., 2010, Winter, 2019). By constraining our model with closed stomata in both the light and dark periods, the model predicted the operation of the starch/sugar-malate cycle as the most energy efficient way to sustain cellular activities. From an evolutionary perspective, if a plant often experiences conditions that require the closure of stomata throughout day and night, such as long periods of severe drought, the evolution of CAM idling would be advantageous for the plant to stay alive. While the evolution of CAM through CAM cycling seems more likely given its similarities to C_3_, it is not impossible that some lineages could establish the starch/sugar-malate cycle through CAM idling.

### Stomatal conductance as a determinant along the C_3_-CAM continuum

It has been proposed that CAM evolution occurs along a continuum from C_3_ to CAM (Silvera et al., 2010; Bräutigam et al., 2017). Our model analysis showed that by varying the CO_2_ exchange in the light period, as a proxy for stomatal conductance, there existed a C_3_-CAM continuum with gradual metabolic changes along the continuum (Figure 2). The key metabolic changes included the processes in the starch/sugar-malate cycle, the TCA cycle at night, and the chloroplastic and mitochondrial ETCs. The fact that a gradual continuum was predicted to be the most energetically favourable way to adapt to a change in stomatal conductance suggests that the fitness landscape between C_3_ and CAM is a smooth one. Given our results, it is not surprising to see many facultative CAM plants which can easily switch between C_3_ and CAM. Based on our model predictions, it is hypothesised that we could find plants anywhere on the C_3_-CAM continuum. A prime example is CAM cycling which falls within the C_3_-CAM continuum at the point when nocturnal CO_2_ exchange is zero. Given the flexibility shown in facultative CAM plants and our results on the C_3_-CAM continuum, it could be possible to find existing plants or engineer new plants that can switch not only between C_3_ and CAM but also at different points on the continuum depending on the environmental conditions.

## Conclusion

Using a core metabolic model of Arabidopsis, we were able to model the metabolic behaviours of CAM, CAM cycling and CAM idling by changing a few simple constraints on gaseous exchange and phloem export. Our results showed that CAM cycling and CAM idling could potentially be evolutionary intermediates on the path to CAM evolution by establishing an intermediate flux through the starch/sugar-malate cycle. By varying the light period CO_2_ exchange as a proxy for stomatal conductance, the model predicted a continuum from C_3_ to CAM with gradual metabolic changes. Besides the insights gained in CAM evolution, the results from this study are informative to guide engineering efforts aiming to introduce CAM into C_3_ crops by identifying the metabolic changes required to convert C_3_ to CAM. In additional to the starch/sugar-malate cycle involved in CAM photosynthesis, our model showed that the fluxes of other metabolic processes, including the TCA cycle and the mitochondrial ETC, need to be altered from C_3_ to optimise CAM.

## List of Supplementary Data

**Supplementary File S1:** Core metabolic model for simulating C_3_, CAM, CAM cycling and CAM idling in SBML and Excel formats

**Supplementary File S2:** Python scripts for running model simulations

**Supplementary Table S1:** Flux solutions from parsimonious flux balance analysis for C_3_, CAM, CAM cycling and CAM idling

**Supplementary Table S2:** A summary of predicted fluxes of key reactions in central metabolism from parsimonious flux balance analysis for C3, CAM, CAM cycling and CAM idling

**Supplementary Table S3:** Flux solutions from parsimonious flux balance analysis for the C_3_-CAM continuum

**Supplementary Table S4:** Flux ranges from flux variability analysis for C_3_, CAM, CAM cycling and CAM idling

**Supplementary Table S5:** Model flux predictions with different malate decarboxylating enzymes

**Supplementary Table S6:** Model flux predictions with different carbohydrate storage

## Acknowledgements

We thank Yale-NUS College (WBS R-607-265-233-121) for the financial support.

