## Supplementary File S2 for "Metabolic modelling of the C_3_-CAM continuum revealed the establishment of a starch/sugar-malate cycle in CAM evolution": CAM.py

```
#CAM
import scobra
m=scobra.Model("C3_CAM.xls")
import os
sim = 'CAM'
if not os.path.isdir(sim):
    os.mkdir(sim)
sim = sim + '/' + sim
constraints_list = {
                   'Starch_biomass1':(0,0),
                   'Starch_biomass2':(0,0),
                   'NADPH_Dehydrogenase_p1':(0,0),
                   'NADPH_Dehydrogenase_p2':(0,0),
                   'Plastoquinol_Oxidase_p1':(0,0),
                   'Plastoquinol_Oxidase_p2':(0,0),
                   'SUCROSE_v_dielTransfer':(0,0),
                   'FRUCTAN_v_dielTransfer':(0,0),
                   }
m.SetConstraints(constraints_list)

#m.SetConstraints({'Photon_tx1':(200,200)})
m.SetConstraints({'Photon_tx2':(0,0)})
m.SetConstraints({'ATPase_tx1':(8.5,8.5)})
m.SetConstraints({'ATPase_tx2':(8.5,8.5)})
m.SetConstraints({'NADPHoxc_tx1':(8.5/9,8.5/9)})
m.SetConstraints({'NADPHoxc_tx2':(8.5/9,8.5/9)})
m.SetConstraints({'NADPHoxm_tx1':(8.5/9,8.5/9)})
m.SetConstraints({'NADPHoxm_tx2':(8.5/9,8.5/9)})
m.SetConstraints({'NADPHoxp_tx1':(8.5/9,8.5/9)})
m.SetConstraints({'NADPHoxp_tx2':(8.5/9,8.5/9)})
m.SetConstraints({'GLC_tx1':(0,0)})
m.SetConstraints({'GLC_tx2':(0,0)})
m.SetConstraints({'SO4_tx1':(0,None),'SO4_tx2':(0,None),'Pi_tx1':(0,None),'Pi_tx2':(0,None)})
m.SetConstraints({'Biomass_tx1':(0,0),'Biomass_tx2':(0,0)})
m.SetConstraints({'Sucrose_tx1':(0,0),'Sucrose_tx2':(0,0)})
m.SetConstraints({'H_tx1':(None,None),'H_tx2':(None,None)})
m.SetConstraints({'NH4_tx1':(0,0),'NH4_tx2':(0,0)})
m.SetConstraints({'unlProtHYPO_c1':(0,0),'unlProtHYPO_c2':(0,0)})
m.SetConstraints({'H_ic1':(None,None),'H_ic2':(None,None)})

#C3
m.SetConstraints({'diel_biomass':(0.258565369)})
#with CO2 and O2 exchange in the day and night
m.SetConstraints({'CO2_tx1':(0,0)})
m.SetConstraints({'O2_tx1':(0,0)})
m.SetConstraints({'CO2_tx2':(None,None)})
m.SetConstraints({'O2_tx2':(None,None)})
m.SetReacsFixedRatio({"RXN_961_p1":1, "RIBULOSE_BISPHOSPHATE_CARBOXYLASE_RXN_p1":5.15})
m.SetReacsFixedRatio({"RXN_961_p2":1, "RIBULOSE_BISPHOSPHATE_CARBOXYLASE_RXN_p2":3})
#free nitrate
#m.SetReacsFixedRatio({"Nitrate_tx1":3, 'Nitrate_tx2':2})

m.SetObjDirec('Min')
m.SetObjective({'Photon_tx1':1,'Photon_tx2':1})
m.MinFluxSolve()
sol=m.GetSol(AsMtx=True)
sol.ToFile(sim + '.csv')
fva = m.FVA(AsMtx=True)
fva.to_csv(sim + '_FVA.csv')

### ME only
m.SetConstraints({'PEPCARBOXYKIN_RXN_c1':(0,0),'PEPCARBOXYKIN_RXN_c2':(0,0)})
m.MinFluxSolve()
sol=m.GetSol(AsMtx=True)
sol.ToFile(sim + '_ME.csv')
fva = m.FVA(AsMtx=True)
fva.to_csv(sim + '_ME_FVA.csv')
m.SetConstraints({'PEPCARBOXYKIN_RXN_c1':(0,None),'PEPCARBOXYKIN_RXN_c2':(0,None)})

### PEPCK only
m.SetConstraints({'MALIC_NADP_RXN_c1':(0,0),'MALIC_NADP_RXN_c2':(0,0),'MALIC_NADP_RXN_p1':(0,0),'MALIC_NADP_RXN_p2':(0,0),'MALIC_NAD_RXN_m1':(0,0),'MALIC_NAD_RXN_m2':(0,0)})
m.MinFluxSolve()
sol=m.GetSol(AsMtx=True)
sol.ToFile(sim + '_PEPCK.csv')
fva = m.FVA(AsMtx=True)
fva.to_csv(sim + '_PEPCK_FVA.csv')
m.SetConstraints({'MALIC_NADP_RXN_c1':(0,None),'MALIC_NADP_RXN_c2':(0,None),'MALIC_NADP_RXN_p1':(0,None),'MALIC_NADP_RXN_p2':(0,None),'MALIC_NAD_RXN_m1':(0,None),'MALIC_NAD_RXN_m2':(0,None)})

### free carbohydrate storage
m.SetConstraints({'SUCROSE_v_dielTransfer':(None,None)})
m.SetConstraints({'FRUCTAN_v_dielTransfer':(None,None)})
m.MinFluxSolve()
sol=m.GetSol(AsMtx=True)
sol.ToFile(sim + '_free_storage.csv')
fva = m.FVA(AsMtx=True)
fva.to_csv(sim + '_free_storage_FVA.csv')

### sucrose only
m.SetConstraints({'STARCH_p_dielTransfer':(0,0)})
m.SetConstraints({'FRUCTAN_v_dielTransfer':(0,0)})
m.MinFluxSolve()
sol=m.GetSol(AsMtx=True)
sol.ToFile(sim + '_sucrose.csv')
fva = m.FVA(AsMtx=True)
fva.to_csv(sim + '_sucrose_FVA.csv')

### fructan only
m.SetConstraints({'SUCROSE_v_dielTransfer':(0,0)})
m.SetConstraints({'FRUCTAN_v_dielTransfer':(None,None)})
m.MinFluxSolve()
sol=m.GetSol(AsMtx=True)
sol.ToFile(sim + '_fructan.csv')
fva = m.FVA(AsMtx=True)
fva.to_csv(sim + '_fructan_FVA.csv')

### Summary
import pandas as pd
ir = pd.read_csv("Interested_Rxn.csv")

starch = pd.read_csv(sim + ".csv")
starch = starch.rename(columns={starch.columns[0]:"Reactions"})
ir = pd.merge(ir, starch, on=["Reactions"], how="left")
ir = ir.rename(columns={ir.columns[-1]:"Starch"})

pepck = pd.read_csv(sim + "_PEPCK.csv")
pepck = pepck.rename(columns={pepck.columns[0]:"Reactions"})
ir = pd.merge(ir, pepck, on=["Reactions"], how="left")
ir = ir.rename(columns={ir.columns[-1]:"PEPCK"})

me = pd.read_csv(sim + "_ME.csv")
me = me.rename(columns={me.columns[0]:"Reactions"})
ir = pd.merge(ir, me, on=["Reactions"], how="left")
ir = ir.rename(columns={ir.columns[-1]:"ME"})

free = pd.read_csv(sim + "_free_storage.csv")
free = free.rename(columns={free.columns[0]:"Reactions"})
ir = pd.merge(ir, free, on=["Reactions"], how="left")
ir = ir.rename(columns={ir.columns[-1]:"Free"})

sucrose = pd.read_csv(sim + "_sucrose.csv")
sucrose = sucrose.rename(columns={sucrose.columns[0]:"Reactions"})
ir = pd.merge(ir, sucrose, on=["Reactions"], how="left")
ir = ir.rename(columns={ir.columns[-1]:"Sucrose"})

fructan = pd.read_csv(sim + "_fructan.csv")
fructan = fructan.rename(columns={fructan.columns[0]:"Reactions"})
ir = pd.merge(ir, fructan, on=["Reactions"], how="left")
ir = ir.rename(columns={ir.columns[-1]:"Fructan"})

ir = ir.fillna(0)
ir.to_csv(sim + "_Summary.csv",index=False)
```
