## Supplementary File S2 for "Metabolic modelling of the C_3_-CAM continuum revealed the establishment of a starch/sugar-malate cycle in CAM evolution": Plotting_C3_CAM_Scan.py

```
import scobra
import matplotlib.pyplot as plt
import numpy as np
import pandas as pd

#     Loading CSV
scan = pd.read_csv("Scan/Scan.csv")

#     Extracting CO2_tx1 data
CO2_tx1_flux = scan[scan['Reactions'] == 'CO2_tx1'].values[0][1:]

#     1. Starch acc + malate accum
starch_flux = scan[scan['Reactions'] == 'STARCH_p_dielTransfer'].values[0][1:]
malate_flux = scan[scan['Reactions'] == 'MAL_v_dielTransfer'].values[0][1:]

#     2. PEPC + PEPCK
PEPC_day_flux = scan[scan['Reactions'] == 'PEPCARBOX_RXN_c1'].values[0][1:]
PEPC_night_flux = scan[scan['Reactions'] == 'PEPCARBOX_RXN_c2'].values[0][1:]
PEPCK_day_flux = scan[scan['Reactions'] == 'PEPCARBOXYKIN_RXN_c1'].values[0][1:]
ME_day_flux = scan[scan['Reactions'] == 'MALIC_NADP_RXN_c1'].values[0][1:]

decarboxylation_flux = PEPCK_day_flux + ME_day_flux
# NOTE there is no PEPCK flux at night

#     3. Photon_tx
Photon_tx = scan[scan['Reactions'] == 'Photon_tx1'].values[0][1:]

#     4. Plastidic + Mitochondrial ATP synthesis (rmb to *3) during the day
Plastidial_ATP_day_flux = scan[scan['Reactions'] == 'Plastidial_ATP_Synthase_p1'].values[0][1:] * 3 
Mitochondrial_ATP_day_flux = scan[scan['Reactions'] == 'Mitochondrial_ATP_Synthase_m1'].values[0][1:] * 3
#Mitochondrial_ATP_night_flux = scan[scan['Reactions'] == 'Mitochondrial_ATP_Synthase_m2'].values[0][1:] * 3
#combined_flux = Plastidial_ATP_day_flux + Mitochondrial_ATP_day_flux + Mitochondrial_ATP_night_flux 

combined_flux = Plastidial_ATP_day_flux + Mitochondrial_ATP_day_flux

#     5. Contribution of mito in producing ATP (in %)
percentage_mito_ATP = np.array([Mitochondrial_ATP_day_flux[i]  / combined_flux[i]  for i in range(len(Plastidial_ATP_day_flux))]) * 100
#percentage_mito_ATP

#     6. Rubisco carboxylase + oxygenase
Rubisco_carboxylase_flux = scan[scan['Reactions'] == 'RIBULOSE_BISPHOSPHATE_CARBOXYLASE_RXN_p1'].values[0][1:]
Rubisco_oxygenase_flux = scan[scan['Reactions'] == 'RXN_961_p1'].values[0][1:]

#     7. Succ deh + NADH dh at night
Succinate_dh_night_flux = scan[scan['Reactions'] == 'SUCCINATE_DEHYDROGENASE_UBIQUINONE_RXN_mi2'].values[0][1:]
NADH_dh_night_flux = scan[scan['Reactions'] == 'NADH_DEHYDROG_A_RXN_mi2'].values[0][1:]

#     8. G6pdh at night
G6P_dh_c_night_flux = scan[scan['Reactions'] == 'GLU6PDEHYDROG_RXN_c2'].values[0][1:]
G6P_dh_p_night_flux = scan[scan['Reactions'] == 'GLU6PDEHYDROG_RXN_p2'].values[0][1:]


#     Creating the figure
from matplotlib.lines import Line2D
from mpl_toolkits.axes_grid1 import make_axes_locatable

fig, ax = plt.subplots(4, 2, figsize=(8,8))
fig.subplots_adjust(top=2.5, right=2, hspace=.35, wspace=.3)

# fonts
axes_fontsize = 20
index_fontsize = 28
xlab_fontsize = 18
ylab_fontsize = 18
legend_fontsize = 18

# colour codes can be found at https://matplotlib.org/examples/color/named_colors.html
colour1='midnightblue'
colour2='sandybrown'
# shape codes can be found at https://matplotlib.org/3.1.0/api/markers_api.html#module-matplotlib.markers
shape1='o'
shape2='s'
# unit
unit = r'$(\mu mol\; m^{-2}\;s^{-1})$'

# 1. Starch acc + malate accum
ax[0,0].scatter(CO2_tx1_flux, starch_flux,s=10,c=colour1, marker=shape1)
ax[0,0].scatter(CO2_tx1_flux, malate_flux,s=10,c=colour2, marker=shape2)
legend_elements = [Line2D([0], [0], color=colour1, marker=shape1, label='Starch'),
                   Line2D([0], [0], color=colour2, marker=shape2, label='Malate')]
ax[0,0].legend(handles=legend_elements, prop={'size': legend_fontsize})
ax[0,0].set_xlim(13.5,0)
ax[0,0].set_xlabel('Light period $\mathregular{CO_2}$ intake ' + unit, fontsize=axes_fontsize)
ax[0,0].set_ylabel('Metabolite accumulation \n' + unit, fontsize=axes_fontsize)
ax[0,0].tick_params(axis="x", labelsize=xlab_fontsize)
ax[0,0].tick_params(axis="y", labelsize=ylab_fontsize)
ax[0,0].text(-0.2, 1.1, 'A', transform=ax[0,0].transAxes, size=index_fontsize, weight='bold')
#ax[0,0].spines['bottom'].set_position('center')
ax[0,0].axhline(linewidth=0.75, color='k')

# 2. PEPC + PEPCK + ME
ax[0,1].scatter(CO2_tx1_flux, PEPC_day_flux + PEPC_night_flux,s=10,c=colour1, marker=shape1)
ax[0,1].scatter(CO2_tx1_flux, decarboxylation_flux,s=10,c=colour2, marker=shape2)
legend_elements = [Line2D([0], [0], color=colour1, marker=shape1, label='PEPC (dark)'),
                   Line2D([0], [0], color=colour2, marker=shape2, label='PEPCK  + ME (light)')]
ax[0,1].legend(handles=legend_elements, prop={'size': legend_fontsize})
ax[0,1].set_xlim(13.5,0)
ax[0,1].set_xlabel('Light period $\mathregular{CO_2}$ intake ' + unit, fontsize=axes_fontsize)
ax[0,1].set_ylabel('PEPC and PEPCK + ME fluxes\n' + unit, fontsize=axes_fontsize)
ax[0,1].tick_params(axis="x", labelsize=xlab_fontsize)
ax[0,1].tick_params(axis="y", labelsize=ylab_fontsize)
ax[0,1].text(-0.2, 1.1, 'B', transform=ax[0,1].transAxes, size=index_fontsize, weight='bold')

# 3. Photon_tx
subsub = ax[1,0]
divider = make_axes_locatable(subsub)
subsub2 = divider.new_vertical(size="1000%", pad=0.1)
fig.add_axes(subsub2)

## bottom partition
subsub.scatter(CO2_tx1_flux, Photon_tx,s=10,c=colour1, marker=shape1)
subsub.set_xlim(13.5,0)
subsub.set_ylim(0, 1)
subsub.tick_params(axis="x", labelsize=xlab_fontsize)
subsub.tick_params(axis="y", labelsize=ylab_fontsize)
yticks = subsub.yaxis.get_major_ticks()
yticks[1].label1.set_visible(False)
subsub.spines['top'].set_visible(False)
subsub.set_xlabel('Light period $\mathregular{CO_2}$ intake ' + unit, fontsize=axes_fontsize)

## top partition 
subsub2.scatter(CO2_tx1_flux, Photon_tx,s=10,c=colour1, marker=shape1)
subsub2.set_xlim(13.5,0)
subsub2.set_ylim(196, 215)
subsub2.tick_params(axis="y", labelsize=ylab_fontsize)
subsub2.tick_params(bottom="off", labelbottom='off')
subsub2.spines['bottom'].set_visible(False)
subsub2.set_ylabel('Photon influx \n' + unit, fontsize=axes_fontsize)
subsub2.text(-0.2, 12.5, 'C', transform=ax[1,0].transAxes, size=index_fontsize, weight='bold')

## From https://matplotlib.org/examples/pylab_examples/broken_axis.html
d = .01
kwargs = dict(transform=subsub2.transAxes, color='k', clip_on=False)
subsub2.plot((-d, +d), (-d, +d), **kwargs)        # top-left diagonal
subsub2.plot((1 - d, 1 + d), (-d, +d), **kwargs)  # top-right diagonal

kwargs.update(transform=subsub.transAxes)  # switch to the bottom axes
subsub.plot((-d, +d), (1 - d*10 , 1 + d*10), **kwargs)  # bottom-left diagonal
subsub.plot((1 - d, 1 + d), (1 - d*10, 1 + d*10), **kwargs)  # bottom-right diagonal

# 4. Plastidial + Mitochondrial ATP synthesis (rmb to *3) during the day
ax[1,1].scatter(CO2_tx1_flux,Plastidial_ATP_day_flux,s=10,c=colour1, marker=shape1)
#ax[1,1].scatter(CO2_tx1_flux,Mitochondrial_ATP_day_flux + Mitochondrial_ATP_night_flux,s=10,c=colour2, marker=shape2)
ax[1,1].scatter(CO2_tx1_flux,Mitochondrial_ATP_day_flux,s=10,c=colour2, marker=shape2)
legend_elements = [Line2D([0], [0], color=colour1, marker=shape1, label='Plastidial ATP synthase'),
                   Line2D([0], [0], color=colour2, marker=shape2, label='Mitochondrial ATP synthase')]
ax[1,1].legend(handles=legend_elements, prop={'size': legend_fontsize})
ax[1,1].set_xlim(13.5,0)
ax[1,1].set_ylim(0, 72)
ax[1,1].set_xlabel('Light period $\mathregular{CO_2}$ intake ' + unit, fontsize=axes_fontsize)
ax[1,1].set_ylabel('Light period ATP synthesis \n' + unit, fontsize=axes_fontsize)
ax[1,1].tick_params(axis="x", labelsize=xlab_fontsize)
ax[1,1].tick_params(axis="y", labelsize=ylab_fontsize)
ax[1,1].text(-0.2, 1.1, 'D', transform=ax[1,1].transAxes, size=index_fontsize, weight='bold')

# 5. Contribution of mito in producing ATP (in %)
subsub = ax[2,0]
divider = make_axes_locatable(subsub)
subsub2 = divider.new_vertical(size="1000%", pad=0.1)
fig.add_axes(subsub2)

## bottom partition
subsub.scatter(CO2_tx1_flux, percentage_mito_ATP,s=10,c=colour1, marker=shape1)
subsub.set_xlim(13.5,0)
subsub.set_ylim(0, 1)
subsub.tick_params(axis="x", labelsize=xlab_fontsize)
subsub.tick_params(axis="y", labelsize=ylab_fontsize)
yticks = subsub.yaxis.get_major_ticks()
yticks[1].label1.set_visible(False)
subsub.spines['top'].set_visible(False)
subsub.set_xlabel('Light period $\mathregular{CO_2}$ intake ' + unit, fontsize=axes_fontsize)

## top partition 
subsub2.scatter(CO2_tx1_flux, percentage_mito_ATP,s=10,c=colour1, marker=shape1)
subsub2.set_xlim(13.5,0)
subsub2.set_ylim(17, 37)
subsub2.tick_params(axis="y", labelsize=ylab_fontsize)
subsub2.tick_params(bottom="off", labelbottom='off')
subsub2.spines['bottom'].set_visible(False)
subsub2.set_ylabel('Mitochondrial contribution in \n light period ATP synthesis (%)', fontsize=axes_fontsize)
subsub2.text(-0.2, 12.5, 'E', transform=ax[2,0].transAxes, size=index_fontsize, weight='bold')

## From https://matplotlib.org/examples/pylab_examples/broken_axis.html
d = .01
kwargs = dict(transform=subsub2.transAxes, color='k', clip_on=False)
subsub2.plot((-d, +d), (-d, +d), **kwargs)        # top-left diagonal
subsub2.plot((1 - d, 1 + d), (-d, +d), **kwargs)  # top-right diagonal

kwargs.update(transform=subsub.transAxes)  # switch to the bottom axes
subsub.plot((-d, +d), (1 - d*10 , 1 + d*10), **kwargs)  # bottom-left diagonal
subsub.plot((1 - d, 1 + d), (1 - d*10, 1 + d*10), **kwargs)  # bottom-right diagonal

# 6. Rubisco carboxylase + oxygenase
ax[2,1].scatter(CO2_tx1_flux,Rubisco_carboxylase_flux,s=10,c=colour1, marker=shape1)
ax[2,1].scatter(CO2_tx1_flux,Rubisco_oxygenase_flux,s=10,c=colour2, marker=shape2)
legend_elements = [Line2D([0], [0], color=colour1, marker=shape1, label='RuBisCO carboxylase'),
                   Line2D([0], [0], color=colour2, marker=shape2, label='RuBisCO oxygenase')]
ax[2,1].legend(handles=legend_elements, prop={'size': legend_fontsize})
ax[2,1].set_xlim(13.5,0)
ax[2,1].set_ylim(0, 17)
ax[2,1].set_xlabel('Light period $\mathregular{CO_2}$ intake ' + unit, fontsize=axes_fontsize)
ax[2,1].set_ylabel('RuBisCO flux \n ' + unit, fontsize=axes_fontsize)
ax[2,1].tick_params(axis="x", labelsize=xlab_fontsize)
ax[2,1].tick_params(axis="y", labelsize=ylab_fontsize)
ax[2,1].text(-0.2, 1.1, 'F', transform=ax[2,1].transAxes, size=index_fontsize, weight='bold')

# 7. Succ dh + NADH dh at night
ax[3,0].scatter(CO2_tx1_flux,Succinate_dh_night_flux,s=10,c=colour1, marker=shape1)
ax[3,0].scatter(CO2_tx1_flux,NADH_dh_night_flux,s=10,c=colour2, marker=shape2)
legend_elements = [Line2D([0], [0], color=colour1, marker=shape1, label='TCA cycle'),
                   Line2D([0], [0], color=colour2, marker=shape2, label='Mitochondrial ETC')]
ax[3,0].legend(handles=legend_elements, prop={'size': legend_fontsize})
ax[3,0].set_xlim(13.5,0)
ax[3,0].set_ylim(0., 4.)
ax[3,0].set_xlabel('Light period $\mathregular{CO_2}$ intake ' + unit, fontsize=axes_fontsize)
ax[3,0].set_ylabel('Fluxes of dark period TCA cycle\nand mitochondrial ETC ' + unit, fontsize=axes_fontsize)
ax[3,0].tick_params(axis="x", labelsize=xlab_fontsize)
ax[3,0].tick_params(axis="y", labelsize=ylab_fontsize)
ax[3,0].text(-0.2, 1.1, 'G', transform=ax[3,0].transAxes, size=index_fontsize, weight='bold')

# 8. G6P dh at night
combined_flux = G6P_dh_c_night_flux + G6P_dh_p_night_flux

subsub = ax[3,1]
divider = make_axes_locatable(subsub)
subsub2 = divider.new_vertical(size="1000%", pad=0.1)
fig.add_axes(subsub2)

## bottom partition
subsub.scatter(CO2_tx1_flux, combined_flux,s=10,c=colour1, marker=shape1)
subsub.set_xlim(13.5,0)
subsub.set_ylim(0, 0.25)
subsub.tick_params(axis="x", labelsize=xlab_fontsize)
subsub.tick_params(axis="y", labelsize=ylab_fontsize)
yticks = subsub.yaxis.get_major_ticks()
yticks[1].label1.set_visible(False)
subsub.spines['top'].set_visible(False)
subsub.set_xlabel('Light period $\mathregular{CO_2}$ intake ' + unit, fontsize=axes_fontsize)

## top partition 
subsub2.scatter(CO2_tx1_flux, combined_flux,s=10,c=colour1, marker=shape1)
subsub2.set_xlim(13.5,0)
subsub2.set_ylim(.94, 1.2)
subsub2.tick_params(axis="y", labelsize=ylab_fontsize)
subsub2.tick_params(bottom="off", labelbottom='off')
subsub2.spines['bottom'].set_visible(False)
subsub2.set_ylabel('Dark period OPPP flux \n ' + unit, fontsize=axes_fontsize)
subsub2.text(-0.2, 12.5, 'H', transform=ax[3,1].transAxes, size=index_fontsize, weight='bold')

## From https://matplotlib.org/examples/pylab_examples/broken_axis.html
d = .01
kwargs = dict(transform=subsub2.transAxes, color='k', clip_on=False)
subsub2.plot((-d, +d), (-d, +d), **kwargs)        # top-left diagonal
subsub2.plot((1 - d, 1 + d), (-d, +d), **kwargs)  # top-right diagonal

kwargs.update(transform=subsub.transAxes)  # switch to the bottom axes
subsub.plot((-d, +d), (1 - d*10 , 1 + d*10), **kwargs)  # bottom-left diagonal
subsub.plot((1 - d, 1 + d), (1 - d*10, 1 + d*10), **kwargs)  # bottom-right diagonal

fig.savefig('Figure 2.png',bbox_inches = 'tight' ,dpi=300)
#plt.show()
```
